# Disruption to functional networks in neonates with perinatal brain injury predicts motor skills at 8 months

**DOI:** 10.1101/226340

**Authors:** Annika C. Linke, Conor Wild, Leire Zubiaurre-Elorza, Charlotte Herzmann, Hester Duffy, Victor K. Han, David S.C. Lee, Rhodri Cusack

## Abstract

**Objective:** Functional connectivity magnetic resonance imaging (fcMRI) of neonates with perinatal brain injury could improve prediction of motor impairment before symptoms manifest, and establish how early brain organization relates to subsequent development. Methods: This cohort study is the first to describe and quantitatively assess functional brain networks and their relation to later motor skills in neonates with a diverse range of perinatal brain injuries. Infants (n=65, included in final analyses: n=53) were recruited from the neonatal intensive care unit (NICU) and were stratified based on their age at birth (premature vs. term), and on whether neuropathology was diagnosed from structural MRI. Functional brain networks and a measure of disruption to functional connectivity were obtained from 14 minutes of fcMRI acquired during natural sleep at term-equivalent age.

**Results:** Disruption to connectivity of the somatomotor and frontoparietal executive networks predicted motor impairment at 4 and 8 months. This disruption in functional connectivity was not found to be driven by differences between clinical groups, or by any of the specific measures we captured to describe the clinical course.

**Conclusion:** fcMRI was predictive over and above other clinical measures available at discharge from the NICU, including structural MRI. Motor learning was affected by disruption to somatomotor networks, but also frontoparietal executive networks, which supports the functional importance of these networks in early development. Disruption to these two networks might be best addressed by distinct intervention strategies.

## 1. Introduction

Thousands of newborns each year are diagnosed with perinatal brain injury secondary to preterm birth, an underlying genetic disorder, asphyxia or neonatal stroke. In a subset of these infants, neonatal brain injury leads to cognitive and behavioral deficits later in life^1–6^. Predicting which infants will develop these delays is difficult, as problems often only become apparent when infants can be assessed behaviorally. This uncertainty puts considerable stress on parents, and hinders targeted early intervention.

Infants with suspected brain injury are often examined using magnetic resonance imaging (MRI). Unfortunately however, the extent of injury visible with routine structural MRI is not always a reliable predictor of long-term developmental outcome. Identifying disruption to brain function with functional connectivity MRI (fcMRI) promises to provide additional information that could improve prediction. In school-age children and adults born prematurely, for example, functional connectivity is altered compared to that of their healthy peers, and these differences are related to measures of developmental outcome, IQ, and performance in school^7–10^. Functional networks can reliably be identified in healthy term-born neonates^11,12^ and even fetuses^13,14^, and it has been suggested that alterations of functional networks as a consequence of premature birth^15–20^ can already be detected at term-equivalent age with fcMRI. It has not yet been determined how these differences relate to neurodevelopmental outcomes, however. Additionally, neonates with even mild neuropathology visible on anatomical MRI scans have been excluded from these studies. In order to understand whether disruptions of functional brain systems due to perinatal brain injury measured at term-equivalent age (TEA) relate to developmental delays detected at follow-up, we studied a cohort of neonates with a diverse range of neuropathologies representative of the perinatal brain injuries commonly encountered in large North American Neonatal Intensive Care Units (NICUs).

We focused on motor function as the outcome measure of interest since it is frequently impacted by perinatal brain injury, is important to daily living, develops rapidly in the first year, and can be measured by observation. Motor skills were assessed at term-equivalent age, and at 4 and 8 months with standard clinical instruments. Our first hypothesis was that fcMRI at TEA would be predictive of motor impairments, over and above other clinical, diagnostic and neurological measures available. We then examined which brain systems were most critical to motor development in this period. We hypothesized that connectivity of the somatomotor network at TEA would be particularly important for motor development in the first year. We furthermore considered which other networks might be relevant. In adults, frontoparietal executive control networks are critical for motor learning^21^. Neuroimaging has shown that these networks are present at term-equivalent age^22,23^, and show the greatest maturational changes in healthy term-born infants over the first two years^24–28^. It has, subsequently, been proposed that they might play a crucial role in infant learning and development^29^, even though there is little behavioral manifestation of executive control before 5 1/2 months postnatally^30,31^. Our third hypothesis was therefore that the frontoparietal executive network would be important for early motor learning. Lastly, we examined whether differences in functional connectivity at TEA and their relationship to motor skills at 8 months could be explained by stratifying infants by prematurity or presence of perinatal brain injury or by any other demographic factors or clinical course in the NICU.

## 2. Materials and Methods

### 2.1 Cohort and MRI

Infants (n=65) were recruited from the tertiary care NICU at Children’s Hospital (LHSC), London, Canada (see Supplementary Methods for inclusion criteria; demographic and clinical information is summarized in Table 1 and Table e-1). Ethical approval was obtained from the Western University Health Sciences REB, and parents gave informed, written consent. Structural and functional MRI were acquired at term-equivalent age (TEA) on a 1.5T 450W GE scanner during unsedated natural sleep (see Supplementary Methods). Sounds were presented during fMRI but the current study examines brain connectivity rather than sound-evoked activation. Functional connectivity has previously been found to be similar between resting-state and tasks including those in which sounds were presented^32^. fMRI images were motion corrected, co-registered to the structural image, and normalized to the UNC neonatal brain template^33^ (see Supplementary Methods for details). Coregistration and normalization were visually inspected (Figure e-1) and 12 datasets were excluded from subsequent analyses due to excessive motion or poor coregistration or normalization.”

**Figure 1.**
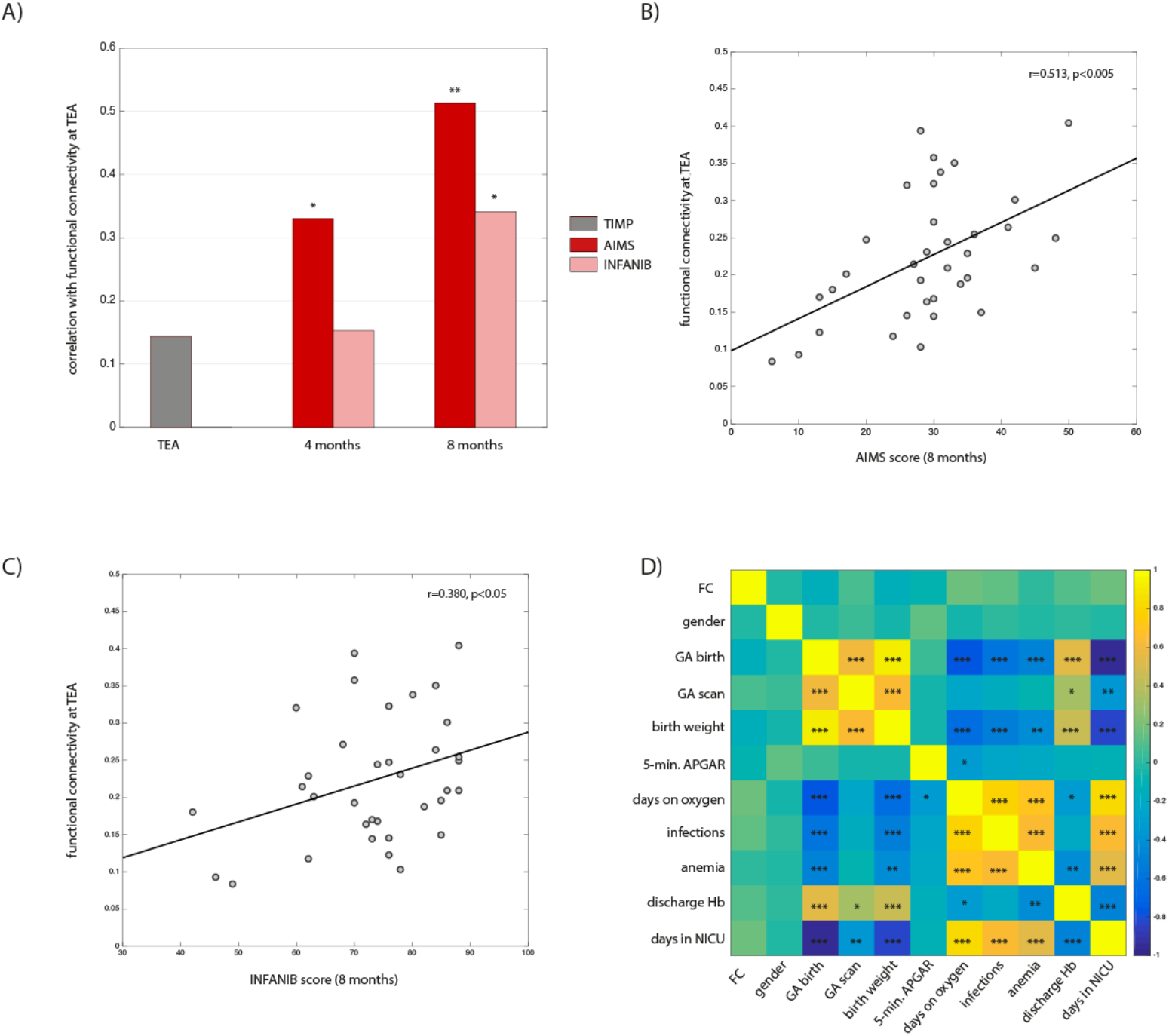
Functional connectivity at term-equivalent age predicted motor skills at 4 and 8 months. (A) Relationship between functional connectivity (FC) at term-equivalent age (TEA) and neurodevelopmental outcome. Correlation scatter plots between functional connectivity and outcome at 8 months are shown in (B) for the AIMS, and (C) for the INFANIB. (D) Relationship between functional connectivity, demographic and clinical information (Pearson correlations). (*** p<0.001, ** p<0.01, * p<0.05)

**Table 1.**
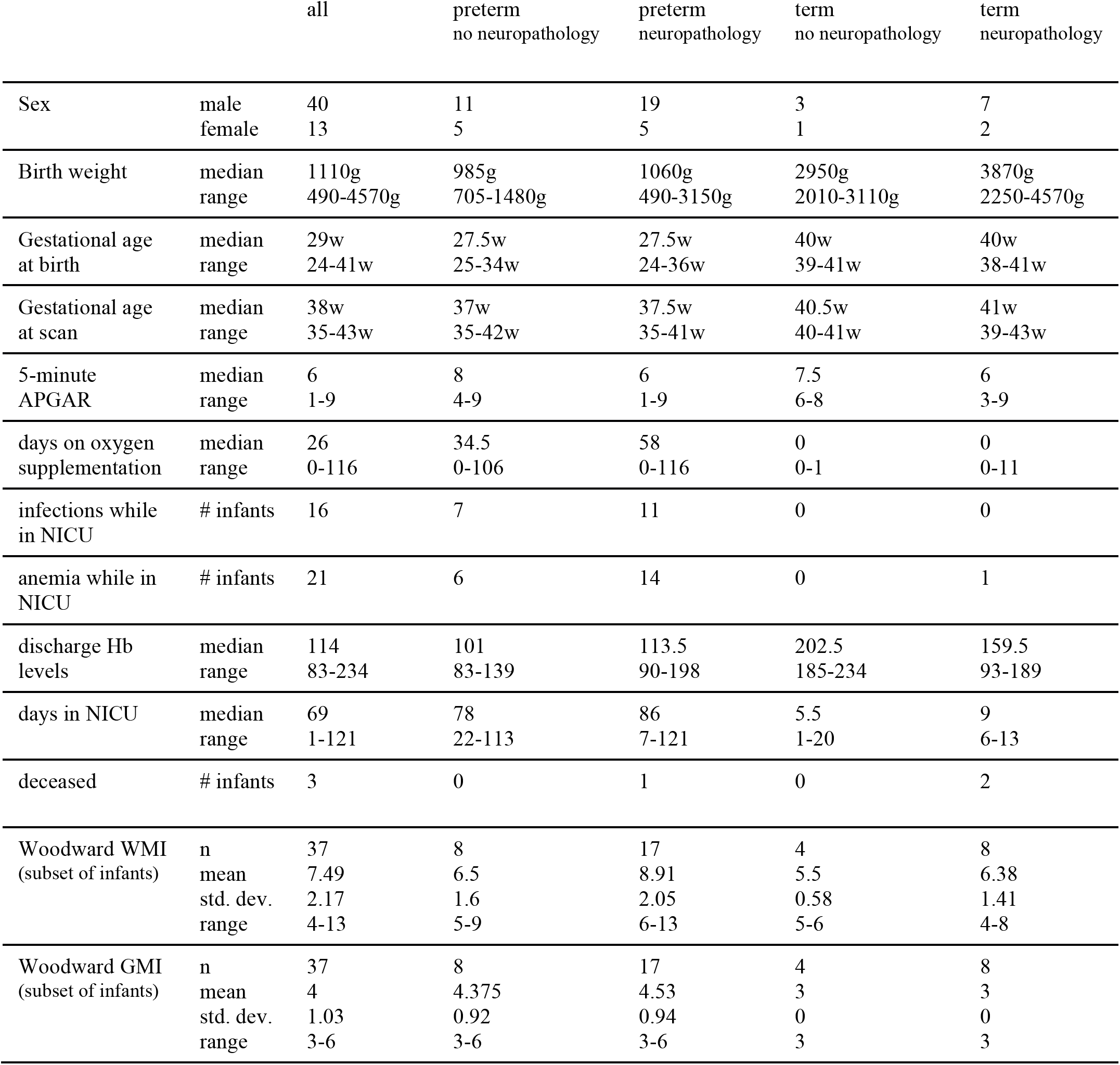
Demographic and Clinical Information.

|  |  | all | preterm<br>no neuropathology | preterm<br>neuropathology | term<br>no neuropathology | term<br>neuropathology |
| --- | --- | --- | --- | --- | --- | --- |
| Sex | male | 40 | 11 | 19 | 3 | 7 |
|  | female | 13 | 5 | 5 | 1 | 2 |
| Birth weight | median | 1110g | 985g | 1060g | 2950g | 3870g |
|  | range | 490-4570g | 705-1480g | 490-3150g | 2010-3110g | 2250-4570g |
| Gestational age<br>at birth | median | 29w | 27.5w | 27.5w | 40w | 40w |
|  | range | 24-41w | 25-34w | 24-36w | 39-41w | 38-41w |
| Gestational age<br>at scan | median | 38w | 37w | 37.5w | 40.5w | 41w |
|  | range | 35-43w | 35-42w | 35-41w | 40-41w | 39-43w |
| 5-minute<br>APGAR | median | 6 | 8 | 6 | 7.5 | 6 |
|  | range | 1-9 | 4-9 | 1-9 | 6-8 | 3-9 |
| days on oxygen<br>supplementation | median | 26 | 34.5 | 58 | 0 | 0 |
|  | range | 0-116 | 0-106 | 0-116 | 0-1 | 0-11 |
| infections while<br>in NICU | # infants | 16 | 7 | 11 | 0 | 0 |
| anemia while in<br>NICU | # infants | 21 | 6 | 14 | 0 | 1 |
| discharge Hb<br>levels | median | 114 | 101 | 113.5 | 202.5 | 159.5 |
|  | range | 83-234 | 83-139 | 90-198 | 185-234 | 93-189 |
| days in NICU | median | 69 | 78 | 86 | 5.5 | 9 |
|  | range | 1-121 | 22-113 | 7-121 | 1-20 | 6-13 |
| deceased | # infants | 3 | 0 | 1 | 0 | 2 |
| Woodward WMI<br>(subset of infants) | n | 37 | 8 | 17 | 4 | 8 |
|  | mean | 7.49 | 6.5 | 8.91 | 5.5 | 6.38 |
|  | std. dev. | 2.17 | 1.6 | 2.05 | 0.58 | 1.41 |
|  | range | 4-13 | 5-9 | 6-13 | 5-6 | 4-8 |
| Woodward GMI<br>(subset of infants) | n | 37 | 8 | 17 | 4 | 8 |
|  | mean | 4 | 4.375 | 4.53 | 3 | 3 |
|  | std. dev. | 1.03 | 0.92 | 0.94 | 0 | 0 |
|  | range | 3-6 | 3-6 | 3-6 | 3 | 3 |

### 2.2 Relating Disruption of Functional Connectivity to Neurodevelopmental Outcome

The pattern of functional connectivity across the brain in each infant at term-equivalent age was compared to the corresponding mean adult pattern of connectivity. Functional connectivity was calculated between every pair of 28 regions-of-interest (ROIs), which were derived from MNI coordinates previously identified in healthy, term-born neonates^34^ (Supplementary Methods and Table e-2). Each ROI comprised an 8 mm sphere at these coordinates, and was normalized to the UNC neonatal template^35^. The mean timecourse of BOLD fMRI activity was extracted for each ROI, and functional connectivity calculated as the Pearson correlation between every pair of timecourses, resulting in a 28x28 connectivity matrix. The similarity of each infant’s connectivity pattern to that of a group of 14 adults yielded a measure of “disruption to functional connectivity” for each infant (Supplementary Methods).

We then assessed whether disruption to functional connectivity at term-equivalent age was related to neurodevelopmental outcome. The infants attended visits at the Developmental Follow-Up Clinic of LHSC, starting shortly after discharge, at which outcome was assessed by trained nurses and clinicians using the Test of Infant Motor Performance (TIMP)^36^ in the first month, and the Alberta Infant Motor Scale (AIMS)^37^ and the Infant Neurological International Battery (INFANIB)^41^ at 4 and 8 months. The degree of disruption to functional connectivity of each infant was then correlated with the TIMP, AIMS and INFANIB scores at each follow-up time point (Supplementary Methods).

To identify which parts of the connectivity matrix drove any correlation between disruption to functional connectivity and outcome, we then decomposed the correlation of the connectivity matrix into the z-scored parts between networks *M* and *N* (where *M=N*corresponds to within network connectivity, and *M*≠*N* between network connectivity).

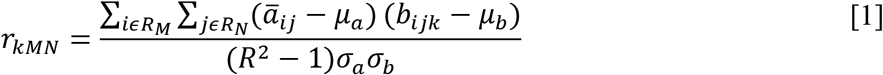

where 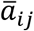 is the mean adult connectivity and *b_ijk_* is the connectivity of infant *k*, between region *i* and region *j*, *R* the total number of regions, *μ_a_* is the mean of the values in the adult matrix and *σ_a_* their standard deviation, *μ_b_*, *σ_b_* the corresponding summary statistics for the infants, and *R_M_* is the set of ROIs within network *M*. Note that the sum of the parts of this decomposition is equal to the original correlation value:

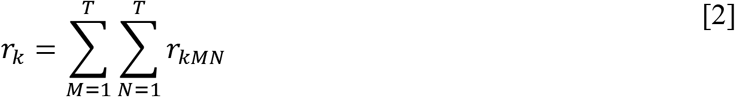

Each of the component measures *r_kMN_* were then taken forwards to a third-order correlation with the outcome, *o_k_*. to yield the contribution of each network separately. We hypothesized that connectivity of the somatomotor network at TEA would be particularly important for motor development in the first year.

### 2.3 Do Differences in Functional Connectivity Reflect Clinical Factors?

Next, we tested whether differences in functional connectivity and their relationship to motor skills at 8 months could be explained by clinical or demographic factors extracted from the NICU discharge reports. We split patients into four pathology groups using two factors each with two levels: preterm vs. term, and presence vs. absence of neuropathology. For each of these four groups, we first established whether five well-established functional brain networks (auditory, visual, motor, default mode and executive control) were equally present in term and preterm infants with perinatal brain injuries. These networks have previously been identified in healthy adults^38^, children^39^, infants and neonates^27,40^, as well as in other patient populations^41,42^, and include regions of the brain involved in a spectrum of functions, from sensory and motor, to higher-level cognition. Group Independent Component Analysis techniques (ICA) ^43^ were adapted for infants with perinatal brain injury using an extension of a method introduced by Wang et al.^44^ - Cross-Validated Regression (CIR) - that avoids circularity and can readily be applied to infant patient populations (see Supplementary Methods). Spatial correlation was used to quantify the similarity of the infants’ functional networks to known network templates^38^. A repeated-measures ANOVA with factor “network” and between-subject factor “pathology group” tested whether the five functional networks could be equally well identified in term and premature infants with and without neuropathology. Similarly, we also tested for any differences in the disruption to functional connectivity measure between the four pathology groups.

Next, we assessed whether disruption of functional connectivity was related to a quantitative scale of brain injury. The Woodward grading system^45^ was applied by a senior neuroradiologist in a subset of infants (n=37, see Supplementary Results). This scoring system grades the degree of perinatal white- and gray-matter injury into four categories: none, mild, moderate and severe. These scores were Pearson correlated with the disruption of functional connectivity measure.

Lastly, we tested whether disruption to functional connectivity was related to specific clinical or demographic factors that might have been lost by stratifying infants into a-priori defined pathology groups. Factors included sex, gestational age at birth and MRI scan, birth weight, 5-minute APGAR scores, days on oxygen supplementation, diagnosis of infections and anemia, discharge hemoglobin levels, and days in the NICU.

## 3. Results

We first tested our central hypothesis, that differences in functional connectivity at term-equivalent age would be related to later motor skills. Results (Figure 1A-C) showed significant positive correlations of individual differences in disruption to functional connectivity measured at TEA with the AIMS (*r*=0.513, *p*<0.005, *CI* [0.217 0.723]) and INFANIB (*r*=0.380, *p*<0.05, *CI* [0.054 0.633]) scores at 8 months. At the same time as behavioral assessments of neurodevelopmental outcome become more reliable^46,47^, correlations become stronger with increasing corrected-age. At 4 months, correlations with the two outcome measures were positive but lower and only significant for the AIMS (*r*=0.330, *p*<0.05, *CI* [0.016 0.585]) but not INFANIB (*r*=0.153, p=0.376, *CI* [-0.180 0.455]) scale. Correlations with the TIMP collected within the first month corrected age were not significant (*r*=0.144, *p*=0.368, *CI* [-0.171 0.433]).

Importantly, while TIMP scores also correlated positively with the AIMS (*r*=0.46, *p*<0.01) and INFANIB (*r*=0.39, *p*<0.05) scores at 8 months of age, partial correlations of the AIMS and INFANIB score with disruption of functional connectivity (controlling for the TIMP scores) remained significant (AIMS: *r_partial_*=0.55, *p*<0.005 vs. *r*=0.61, *p*<0.001; INFANIB: *r_partial_*=0.46, *p*<0.01 vs. *r*=0.53, *p*<0.005 for the 31 infants with both TIMP and AIMS/INFANIB scores).

Given the predictive value of disruption to functional connectivity, we assessed which of seven functional networks among the 28 ROIs^34^ (language-LAN, sensorimotor-SMN, visual-VIS, default mode-DMN, dorsal attention-DAN, ventral attention-VAN and frontoparietal control-FPC) drove the correlation of whole-brain functional connectivity patterns with neurodevelopmental outcome most strongly. Given that we focused on motor outcome, we predicted connectivity of the motor network at TEA to considerably influence infant motor development. Additionally, the frontoparietal executive network is crucial for learning in later life but its role in early infant development is not known. We predicted that even before first behavioral signs of executive function emerge, this network already plays an important role for skill learning, including motor development. This was indeed what we found (Table 2). Our results show that connectivity within the SMN and between the SMN-DMN and SMN-VAN contributed most to the correlation with motor skills at 8 months. Additional connectivity within the FPC and between the DMN and VIS also contributed.”

**Table 2.**
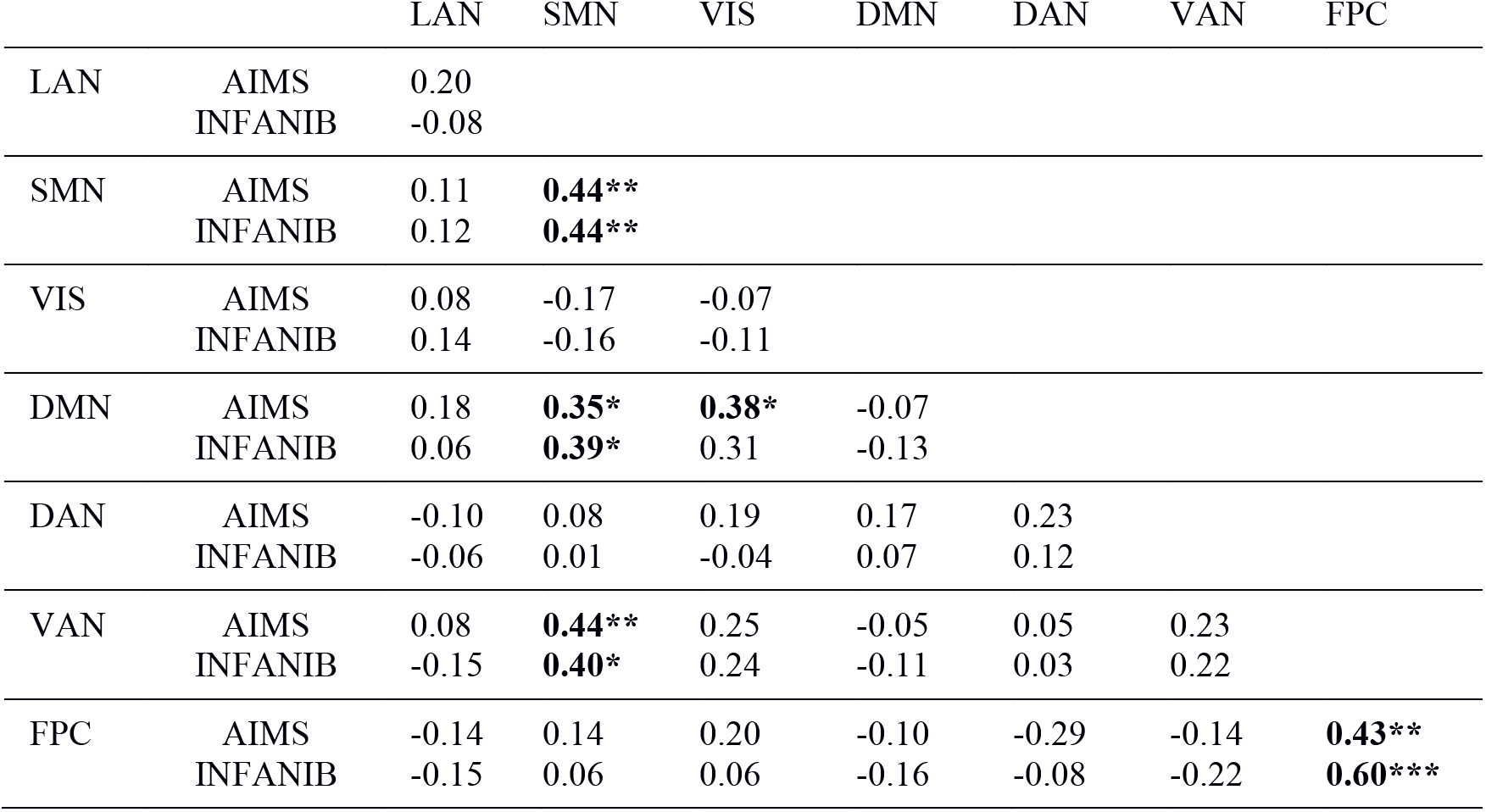
Networks driving correlation of functional connectivity with outcome at 8 months (values are correlation coefficient rho, * indicates significance at *p*<0.05, ** p<0.01, ***p<0.001)

We then tested whether differences in functional connectivity were related to demographic and clinical factors by, first, stratifying infants by prematurity (preterm/term) and presence/absence of neuropathology. Visual inspection and a repeated-measures ANOVA suggested the infants’functional networks were similar to normative template networks^38^ in healthy adults irrespective of pathology group (F(3,49)=0.653, p=0.585, η^2^=0.038, Figure 2 and Figure 3A, also see Supplementary Results). Similarly, disruption to functional connectivity was not found to be different between groups (*F*(3,49)=0.226, *p*=0.87S, *η*2=0.014, Figure 3B, Figure e-3).

**Figure 2.**
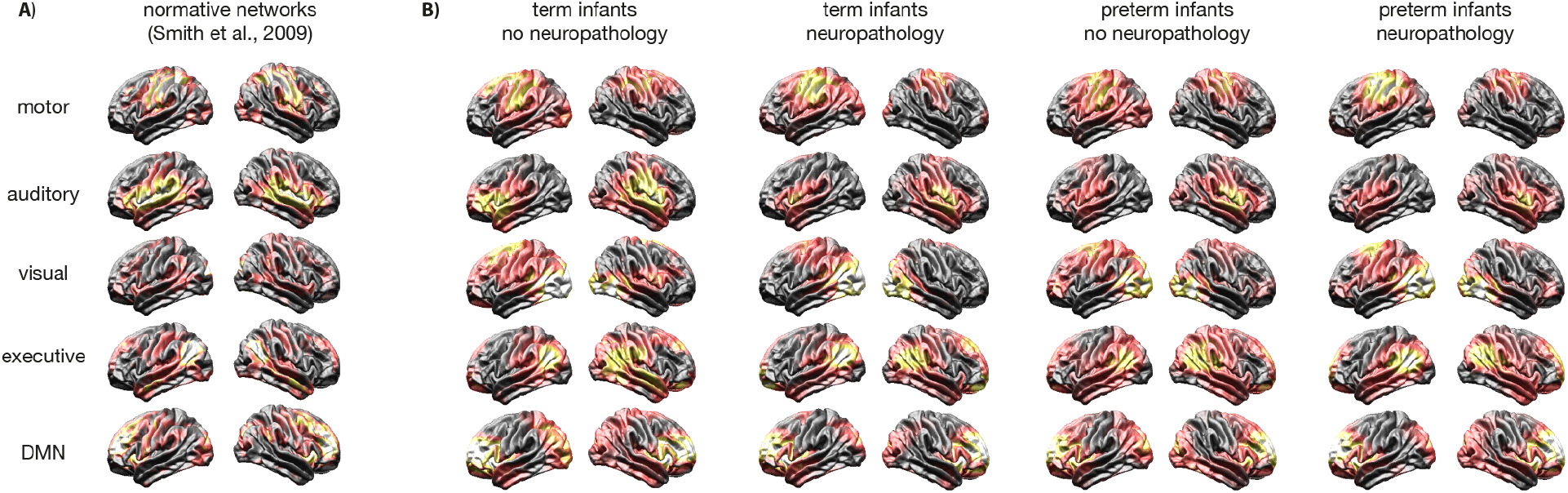
Corresponding functional networks in adults and infants. Functional networks (A) in healthy adults (Smith et al., 2009) that were used as templates during Cross-Iterative Regression (CIR), and (B) as derived in infants, split by pathology group. Lighter colors indicate stronger evidence of the respective network. Spatial topography of each network was similar to the adult templates in all four infant pathology groups.

**Figure 3.**
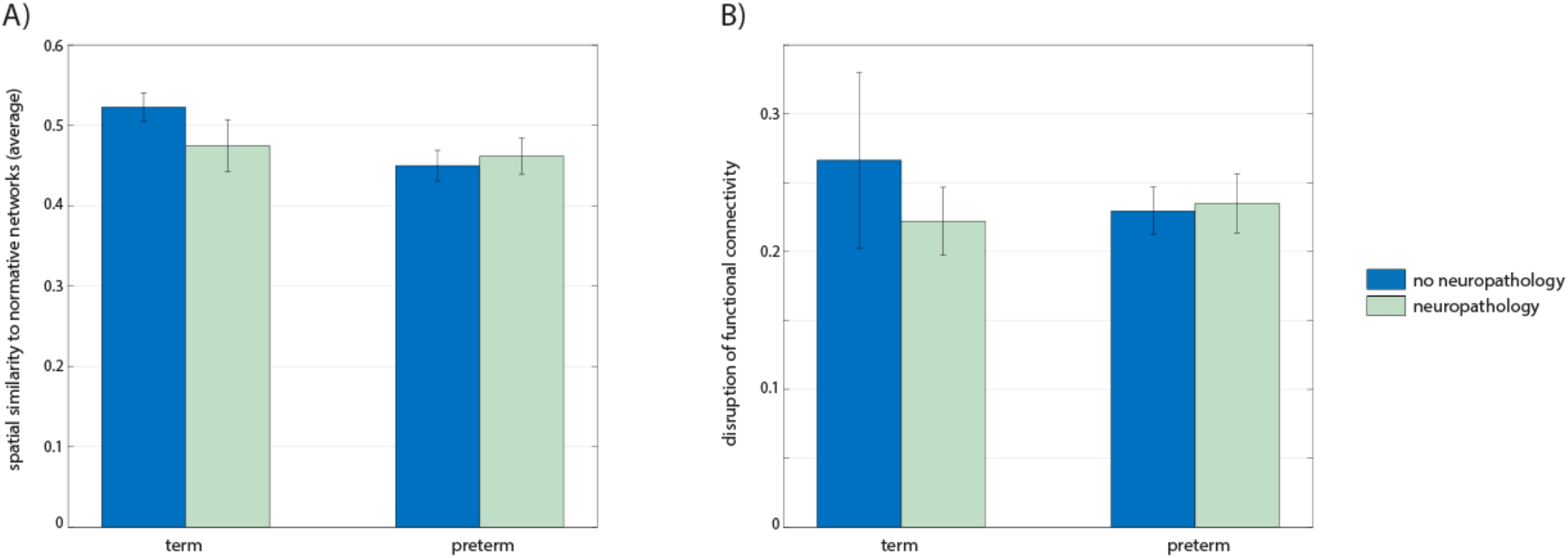
Functional connectivity did not differ consistently between groups. No significant differences between the four infant pathology groups in (A) functional network topography (CIR analysis, average of all networks shown), and (B) patterns of functional connectivity. Error bars are standard errors.

The presence or absence of neuropathology is a crude measure of the degree of brain injury. The Woodward grading system was therefore used to quantify the degree of brain injury in a subset of infants (n=37, also see Supplementary Results). Functional connectivity was not related to this quantitative measure of brain injury, for neither white-matter (*r*=0.04, *p*=0.814, *CI* [-0.288 0.359]) nor grey-matter (*r*=-0.101, *p*=0.552, *CI* [-0.411 0.231]).

Importantly, adverse outcome is typically only observed in a subset of NICU infants^48–50^. Grouping infants into heterogeneous categories such as premature/term birth and presence of neuropathology might decrease sensitivity to detect subtle alterations in functional connectivity that are related to later differences in behavior. We, lastly, tested whether ten clinical or demographic factors that might not have been captured by the four pathology groups were related to disruption to functional connectivity. While many of the clinical variables are highly correlated (Figure 1D), differences in disruption to functional connectivity were not significantly related to any of them (top row). Additionally, clinical and demographic factors were not related to motor skills at 8 months as assessed by the TIMP, AIMS and INFANIB (Table e-3). These results likely reflect the difficulty of predicting neurodevelopmental outcome from clinical information available at discharge from the NICU. Our results suggest that functional connectivity measured at term-equivalent age provides additional information that is independent from currently available clinical information, and that can contribute to the prediction of neurodevelopmental outcome after preterm birth and perinatal brain injury.

## 4. Discussion

This study shows that it is possible to robustly identify functional brain networks in infants with perinatal brain injuries at TEA, paving the way for future studies of this vulnerable clinical population. Differences in functional connectivity irrespective of pathology group correlated significantly with motor skills at 4 and 8 months. Specifically, disruption to the motor and frontoparietal executive networks drove this relationship most strongly. This implies that fMRI provides prognostic information at the time of discharge from the NICU. Our results extend previous findings by Arichi et al.^51^ who found substantial motor network connectivity abnormalities in three neonates with severe hemorrhagic parenchymal infarction who later developed cerebral palsy (CP). Similarly, a study in 14 infants with moderate to severe white matter injury secondary to periventricular hemorrhagic infarction^52^ found that functional connectivity was disrupted, particularly in the motor network and cerebellar regions. Compared to these two studies, however, most infants in the current cohort had milder and more diverse neuropathologies, and including cortical regions that spanned seven distinct functional networks allowed us to assess the relationship between motor outcome and brain function across cortex. This is important, as the most common perinatal brain injuries (such as low-grade intraventricular hemorrhage following premature birth) put an infant at increased risk of developmental delays that are much harder to detect early than CP.

Our results suggest that the executive system may be important for development much earlier than previously thought^29^. Injury to this system essential for learning and cognition would be expected to lead to a spectrum of neurodevelopmental deficits. Smyser et al.^34^ also found alterations in functional connectivity of the DMN and FPC in premature infants scanned at term-equivalent age, but since no behavioural follow-up information was included, it remained unknown whether this influenced development. Importantly, screening tests like the AIMS and INFANIB provide the first signs not only of motor disability but also of more general neurodevelopmental delays and disorders, and it has consequently been argued that all infants should undergo developmental motor screening at the end of the first year^53^. By 8 months, motor milestones are predictive of various aspects of later development, even when controlling for gestational age, birth weight, and disability^54^. Long-term follow-up information would provide important insights into the power of fcMRI collected at term-equivalent age to improve early prediction of broader cognitive and social outcomes for infants at risk.

Our results also show that studying neonatal brain function in predefined groups might miss variability in the data that explains later developmental outcome. A number of previous studies have found functional networks to be surprisingly similar in premature and healthy-term born infants with the subset of networks altered varying greatly between studies^18,19,55–57^, and with some finding no differences at all^25,42^. This seems at odds with the higher incidence of developmental delays in premature infants, and the abnormalities in functional connectivity found in older children and adults born prematurely^9,10,58,59^. It is possible that differences in functional connectivity only emerge over time. Alternatively, these findings might reflect a lack of sensitivity to pick up relatively subtle and diverse differences between groups defined a-priori. Approximately 30% of extremely premature infants will develop moderate or severe developmental delays and disability^48–50^. Since infants with signs of neuropathology were excluded from previous studies investigating functional network maturity and disruption after premature birth, the risk of developmental delays for the infants typically included in neonatal fcMRI studies is likely much lower. As such, it is reassuring that functional brain organization seems to be unaltered in the majority of “healthy” preterm neonates. Those with developmental delays and clear disruptions of functional connectivity, on the other hand, might be over-represented in studies of older children and adults born prematurely.

Three more recent studies employing advanced statistical methods that have more power to detect subtle differences found disruptions of functional connectivity in premature infants without perinatal brain injury^15,34,60^. These studies did not assess whether alterations of functional connectivity predicted developmental outcome. However, another recent study^61^ showed thalamocortical connectivity measured at 1 year correlated with assessments of cognitive function at 2 years (n=143). It is even more important to assess such relationships in infants at high risk of developmental delays, like those born prematurely or those who have sustained perinatal brain injury. The current study is an important step in this direction.

We hope our results will encourage others to study infants with perinatal brain injury using fMRI, so that limitations in the current study can be addressed. Most importantly, the inclusion criteria in our study were broad in order to be able to collect a sample reflecting commonly encountered neuropathologies in North American NICUs. This meant that when grouping infants by age at birth and presence of neuropathology, sample sizes were moderate and unbalanced. Some differences between these groups might become significant with larger sample sizes. However, neither demographics, the clinical course in the NICU, or the Woodward grading of the degree brain injury (irrespective of a-priori group membership) were related to network connectivity.

Perinatal brain injury is common in NICU infants but early prediction of outcome is difficult, leading to delays in interventions, increased medical expenditures and anxiety and stress for parents and caregivers. Functional MRI may offer valuable independent information to aid the prediction of neurodevelopmental outcome at TEA irrespective of the clinical course in the NICU or the brain injury acquired. We hope that this will facilitate earlier, focused intervention to address the functional disruption in infants, and decrease the uncertainty parents currently face.

## Abbreviations

CIR: – Cross-validated Iterative Regression
fMRI: – functional Magnetic Resonance Imaging
EPI: – Echo Planar Imaging
ICA: – Independent Component Analysis
IVH: – Intraventricular Hemorrhage

## Acknowledgements

We thank the families of the infants in this study, the NICU nurses and the MRI technicians at Children’s Hospital (LHSC), London, Ontario, Canada, and Richa Metha, Andrea Lum and Keng Yeow Tay for their continued enthusiasm, patience, and support. We thank Deborah Ness for her assistance in formatting this manuscript.

## References

1 van Buuren LM, van der Aa NE, Dekker HC, et al. Cognitive outcome in childhood after unilateral perinatal brain injury. Dev Med Child Neurol. 2013;55(10):934–940. doi:10.1111/dmcn.12187.

2 Farooqi A, Hägglöf B, Sedin G, Serenius F. Impact at age 11 years of major neonatal morbidities in children born extremely preterm. Pediatrics. 2011;127(5):e1247–57. doi:10.1542/peds.2010-0806.

3 Miller SP, Ramaswamy V, Michelson D, et al. Patterns of brain injury in term neonatal encephalopathy. J Pediatr. 2005;146(4):453–460. doi:10.1016/j.jpeds.2004.12.026.

4 Peterson BS. Regional Brain Volume Abnormalities and Long-term Cognitive Outcome in Preterm Infants. JAMA. 2000;284(15):1939. doi:10.1001/jama.284.15.1939.

5 Hack M. Perinatal Brain Injury in Preterm Infants and Later Neurobehavioral Function. JAMA. 2000;284(15):1973. doi:10.1001/jama.284.15.1973.

6 Inder TE. Pediatrics: Predicting outcomes after perinatal brain injury. Nat Rev Neurol. 2011;7(10):544–545. doi:10.1038/nrneurol.2011.142.

7 Damaraju E, Phillips JR, Lowe JR, Ohls R, Calhoun VD, Caprihan A. Resting-state functional connectivity differences in premature children. Front Syst Neurosci. 2010;4(June):1–13. doi:10.3389/fnsys.2010.00023.

8 Gozzo Y, Vohr B, Lacadie C, et al. Alterations in neural connectivity in preterm children at school age. Neuroimage. 2009;48(2):458–463. doi:10.1016/j.neuroimage.2009.06.046.

9 Dick AS, Raja Beharelle A, Solodkin A, Small SL. Interhemispheric Functional Connectivity following Prenatal or Perinatal Brain Injury Predicts Receptive Language Outcome. J Neurosci. 2013;33(13):5612–5625. doi:10.1523/JNEUROSCI.2851-12.2013.

10 Schafer RJ, Lacadie C, Vohr B, et al. Alterations in functional connectivity for language in prematurely born adolescents. Brain. 2009;132(Pt 3): 661–670. doi:10.1093/brain/awn353.

11 van den Heuvel MP, Kersbergen KJ, de Reus MA, et al. The Neonatal Connectome During Preterm Brain Development. Cereb Cortex. May 2014: 1–14. doi:10.1093/cercor/bhu095.

12 Fransson P, Skiöld B, Engström M, et al. Spontaneous brain activity in the newborn brain during natural sleep--an fMRI study in infants born at full term. Pediatr Res. 2009;66(3): 301–305. doi:10.1203/PDR.0b013e3181b1bd84.

13 Thomason ME, Grove LE, Lozon TA, et al. Age-related increases in long-range connectivity in fetal functional neural connectivity networks in utero. Dev Cogn Neurosci. September 2014. http://www.sciencedirect.com/science/article/pii/S1878929314000644. Accessed January 15, 2015.

14 Thomason ME, Dassanayake MT, Shen S, et al. Cross-hemispheric functional connectivity in the human fetal brain. Sci Transl Med. 2013; 5(173): 173ra24. http://www.pubmedcentral.nih.gov/articlerender.fcgi?artid=3618956&tool=pmcentrez&rendertype=abstract. Accessed January 15, 2015.

15 Ball G, Aljabar P, Arichi T, et al. Machine-learning to characterise neonatal functional connectivity in the preterm brain. Neuroimage. 2016;124:267–275. doi:10.1016/j.neuroimage.2015.08.055.

16 Smyser CD, Snyder AZ, Shimony JS, Mitra A, Inder TE, Neil JJ. Resting-State Network Complexity and Magnitude Are Reduced in Prematurely Born Infants. Cereb Cortex. October 2014:1–12. doi:10.1093/cercor/bhu251.

17 Scheinost D, Kwon SH, Shen X, et al. Preterm birth alters neonatal, functional rich club organization. Brain Struct Funct. 2015. doi:10.1007/s00429-015-1096-6.

18 Kwon SH, Scheinost D, Lacadie C, et al. Adaptive Mechanisms of Developing Brain: Cerebral Lateralization in the Prematurely-Born. Neuroimage. 2015;108:144–150.

19 Toulmin H, Beckmann CF, O’Muircheartaigh J, et al. Specialization and integration of functional thalamocortical connectivity in the human infant. Proc Natl Acad Sci. 2015;112(20):6485–6490. doi:10.1073/pnas.1422638112.

20 Lin W, Zhu Q, Gao W, et al. Functional connectivity MR imaging reveals cortical functional connectivity in the developing brain. AJNR Am J Neuroradiol. 2008;29(10):1883–1889. doi:10.3174/ajnr.A1256.

21 Miller EK, Cohen JD. An integrative theory of prefrontal cortex function. Annu Rev Neurosci. 2001;24(1):167–202. doi:10.1146/annurev.neuro.24.1.167.

22 van den Heuvel MP, Kersbergen KJ, de Reus MA, et al. The Neonatal Connectome During Preterm Brain Development. Cereb Cortex. May 2014:1–14. doi:10.1093/cercor/bhu095.

23 Fransson P, Skiöld B, Engström M, et al. Spontaneous brain activity in the newborn brain during natural sleep-an fMRI study in infants born at full term. Pediatr Res. 2009;66(3):301–305. doi:10.1203/PDR.0b013e3181b1bd84.

24 Gao W, Alcauter S, Smith JK, Gilmore JH, Lin W. Development of human brain cortical network architecture during infancy. Brain Struct Funct. 2015;220:1173–1186. doi:10.1007/s00429-014-0710-3.

25 Doria V, Beckmann CF, Arichi T, et al. Emergence of resting state networks in the preterm human brain. Proc Natl Acad Sci U S A. 2010;107(46):20015–20020. doi:10.1073/pnas.1007921107.

26 Cao M, He Y, Dai Z, et al. Early Development of Functional Network Segregation Revealed by Connectomic Analysis of the Preterm Human Brain. Cereb Cortex. 2016:bhw038. doi:10.1093/cercor/bhw038.

27 Fransson P, Skiöld B, Horsch S, et al. Resting-state networks in the infant brain. Proc Natl Acad Sci U S A. 2007;104(39):15531–15536. doi:10.1073/pnas.0704380104.

28 Fransson P, Aden U, Blennow M, Lagercrantz H. The functional architecture of the infant brain as revealed by resting-state fMRI. Cereb Cortex. 2011;21(1):145–154. doi:10.1093/cercor/bhq071.

29 Cusack R, Ball G, Smyser CD, Dehaene-Lambertz G. A Neural Window on the Emergence of Cognition. Ann N Y Acad Sci. 2016;1369(1):1–18. doi:10.1111/nyas.13036.

30 Reznick JS, Morrow JD, Goldman BD, Snyder J. The onset of working memory in infants. Infancy. 2004;6(1):145–154. doi:10.1207/s15327078in0601_7.

31 Reynolds GD, Romano AC. The Development of Attention Systems and Working Memory in Infancy. Front Syst Neurosci. 2016; 10:Article 15. doi:10.3389/fnsys.2016.00015.

32 Shah LM, Cramer JA, Ferguson MA, Birn RM, Anderson JS. Reliability and reproducibility of individual differences in functional connectivity acquired during task.and resting state. Brain Behav. 2016;6(5):e00456. doi:10.1002/brb3.456.

33 Shi F, Yap P-T, Wu G, et al. Infant brain atlases from neonates to 1- and 2-year-olds. PLoS One. 2011;6(4):e18746. doi:10.1371/journal.pone.0018746.

34 Smyser CD, Snyder AZ, Shimony JS, Mitra A, Inder TE, Neil JJ. Resting-State Network Complexity and Magnitude Are Reduced in Prematurely Born Infants. Cereb Cortex. October 2014:1–12. doi:10.1093/cercor/bhu251.

35 Shi F, Yap P-T, Wu G, et al. Infant brain atlases from neonates to 1- and 2-year-olds. PLoS One. 2011;6(4):e18746. doi:10.1371/journal.pone.0018746.

36 Campbell SK, Kolobe TH, Osten ET, Lenke M, Girolami GL. Construct validity of the test of infant motor performance. Phys Ther. 1995;75(7):585–596. http://www.ncbi.nlm.nih.gov/pubmed/7604077. Accessed March 22, 2016.

37 Piper MC, Pinnell LE, Darrah J, Maguire T, Byrne PJ. Construction and validation of the Alberta Infant Motor Scale (AIMS). Can J Public Heal. 1992;83:46–50. http://www.ncbi.nlm.nih.gov/pubmed/1468050. Accessed March 22, 2016.

38 Smith SM, Fox PT, Miller KL, et al. Correspondence of the brain’s functional architecture during activation and rest. PNAS. 2009;106(31):13040–13045.

39 de Bie HMA, Boersma M, Adriaanse S, et al. Resting-state networks in awake five-to eight-year old children. Hum Brain Mapp. 2012;33(5):1189–1201. doi:10.1002/hbm.21280.

40 Gao W, Alcauter S, Elton A, et al. Functional Network Development During the First Year: Relative Sequence and Socioeconomic Correlations. Cereb Cortex. 2015;25:291902928. doi:10.1093/cercor/bhu088.

41 Greicius MD, Kiviniemi V, Tervonen O, Vainionpää V, Reiss AL, Menon V. Persistent Default-Mode Network Connectivity During Light Sedation. Hum Brain Mapp. 2008;29(7):839–847. doi:10.1002/hbm.20537.Persistent.

42 Lee W, Morgan BR, Shroff MM, Sled JG, Taylor MJ. The development of regional functional connectivity in preterm infants into early childhood. Neuroradiology. 2013;55 Suppl 2:105–111. doi:10.1007/s00234-013-1232-z.

43 Calhoun VD, Liu J, Adali T. A review of group ICA for fMRI data and ICA for joint inference of imaging, genetic, and ERP data. Neuroimage. 2009;45(1 Suppl):S163–72. doi:10.1016/j.neuroimage.2008.10.057.

44 Wang D, Buckner RL, Fox MD, et al. Parcellating cortical functional networks in individuals. Nat Neurosci. 2015;18(12):1853–1860. doi:10.1038/nn.4164.

45 Woodward LJ, Anderson PJ, Austin NC, Howard K, Inder TE. Neonatal MRI to Predict Neurodevelopmental Outcomes in Preterm Infants. N Engl J Med. 2006;355(7):685–694.

46 Pedersen S, Sommerfelt K, Markestad T. Early motor development of premature infants with birthweight less than 2000 grams. Acta Paediatr. 2007;89(12):1456–1461. doi:10.1111/j.1651-2227.2000.tb02776.x.

47 Campbell SK, Hedeker D. Validity of the Test of Infant Motor Performance for discriminating among infants with varying risk for poor motor outcome. J Pediatr. 2001;139(4):546–551. doi:10.1067/mpd.2001.117581.

47 Serenius F, Källén K, Blennow M, et al. Neurodevelopmental Outcome in Extremely Preterm Infants at 2.5 Years After Active Perinatal Care in Sweden. JAMA. 2013;309(17):1810. doi:10.1001/jama.2013.3786.

49 Allen MC. Neurodevelopmental outcomes of preterm infants. Curr Opin Neurol.2008;21:123–128.

50 Marlow N, Wolke D, Bracewell MA, Samara M. Neurologic and Developmental Disability at Six Years of Age after Extremely Preterm Birth. N Engl J Med. 2005;352(1):9–19. doi:10.1056/NEJMoa041367.

51 Arichi T, Counsell SJ, Allievi AG, et al. The effects of hemorrhagic parenchymal infarction on the establishment of sensori-motor structural and functional connectivity in early infancy. Neuroradiology. 2014;56(11):985–994. doi:10.1007/s00234-014-1412-5.

52 Smyser CD, Snyder AZ, Shimony JS, Blazey TM, Inder TE, Neil JJ. Effects of White Matter Injury on Resting State fMRI Measures in Prematurely Born Infants. Fan Y, ed. PLoS One. 2013;8(7):e68098. doi:10.1371/journal.pone.0068098.

53 Harris SR. A plea for developmental motor screening in Canadian infants. Paediatr Child Health. 2016;21(3):129–130.

54 Ghassabian A, Sundaram R, Bell E, Bello SC, Kus C, Yeung E. Gross Motor Milestones and Subsequent Development. Pediatrics. 2016.

55 Smyser CD, Snyder AZ, Neil JJ. Functional connectivity MRI in infants: exploration of the functional organization of the developing brain. Neuroimage. 2011;56(3):1437–1452. doi:10.1016/j.neuroimage.2011.02.073.

56 Smyser CD, Inder TE, Shimony JS, et al. Longitudinal Analysis of Neural Network Development in Preterm Infants. Cereb Cortex. 2010;20(December):2852–2862. doi:10.1093/cercor/bhq035.

57 Lin W, Zhu Q, Gao W, et al. Functional connectivity MR imaging reveals cortical functional connectivity in the developing brain. AJNR Am J Neuroradiol. 2008;29(10):1883–1889. doi:10.3174/ajnr.A1256.

58 Damaraju E, Phillips JR, Lowe JR, Ohls R, Calhoun VD, Caprihan A. Resting-state functional connectivity differences in premature children. Front Syst Neurosci. 2010;4(June):1–13. doi:10.3389/fnsys.2010.00023.

59 Gozzo Y, Vohr B, Lacadie C, et al. Alterations in neural connectivity in preterm children at school age. Neuroimage. 2009;48(2):458–463. doi:10.1016/j.neuroimage.2009.06.046.

60 Scheinost D, Kwon SH, Shen X, et al. Preterm birth alters neonatal, functional rich club organization. Brain Struct Funct. 2015. doi:10.1007/s00429-015-1096-6.

61 Alcauter S, Lin W, Smith JK, et al. Frequency of spontaneous BOLD signal shifts during infancy and correlates with cognitive performance. Dev Cogn Neurosci. 2014;12C:40–50. doi:10.1016/j.dcn.2014.10.004.

